# Competition in biofilms between cystic fibrosis isolates of *Pseudomonas aeruginosa* is shaped by R-pyocins

**DOI:** 10.1101/264580

**Authors:** Olubukola Oluyombo, Christopher N. Penfold, Stephen P. Diggle

## Abstract

*Pseudomonas aeruginosa* is an opportunistic pathogen responsible for a number of different human infections and is the leading cause of morbidity and mortality in cystic fibrosis (CF) patients. *P. aeruginosa* infections are difficult to treat due to a number of antibiotic resistance mechanisms and the organisms propensity to form multicellular biofilms. Epidemic strains of *P. aeruginosa* often dominate within the lungs of individual CF patients, but how they achieve this is poorly understood. One of the ways strains of *P. aeruginosa* can compete, is by producing chromosomally encoded bacteriocins, called pyocins. Three major classes of pyocin have been identified in *P. aeruginosa:* soluble pyocins (S-types) and tailocins (R- and F-types). In this study, we investigated the distribution of S- and R-type pyocins in 24 clinical strains isolated from individual CF patients and then focused on understanding their roles on inter-strain competition. We found that (i) each strain produced only one R-pyocin type, but the number of S-pyocins varied between strains; (ii) R-pyocins were generally important for strain dominance during competition assays in planktonic cultures and biofilm communities in strains with both disparate R and S pyocin sub-types. (iii) purified R-pyocins demonstrated significant antimicrobial activity against established biofilms. Our work provides support for a key role played by R-pyocins in the competition between *P. aeruginosa* strains, and may help explain why certain strains and lineages of *P. aeruginosa* dominate and displace others during CF lung infection. Furthermore, we demonstrate the potential of exploiting R-pyocins for therapeutic gains in an era when antibiotic resistance is a global concern.

**IMPORTANCE:** A major clinical problem caused by *Pseudomonas aeruginosa*, is chronic biofilm infection of the lungs in individuals with cystic fibrosis (CF). Epidemic *P. aeruginosa* strains dominate and displace others during CF infection, but these intra-species interactions remain poorly understood. Here we demonstrate that R-pyocins (bacterocins) are important factors in driving competitive interactions in biofilms between *P. aeruginosa* strains isolated from different CF patients. In addition, we found that these phage-like pyocins are inhibitory against mature biofilms of susceptible strains. This highlights the potential of R-pyocins as antimicrobial and antibiofilm agents, at a time when new antimicrobial therapies are desperately needed.

## Introduction

*Pseudomonas aeruginosa* is an opportunistic pathogen, capable of infecting different host species, including plants, insects, and mammals (1). It is intrinsically resistant to many classes of antibiotic, and produces a range of tissue damaging extracellular products such as exoenzymes and phenazine pigments which aid dissemination and spread within a host (2). A major clinical problem caused by *P. aeruginosa*, is chronic infection of the lungs in individuals with cystic fibrosis (CF), where it contributes significantly to morbidity over the lifetime of the patient and accelerates mortality (3). Strains that thrive in CF lung infections evolve in the lung environment over time, undergoing genomic mutations and rearrangements which results in phenotypic variation within populations (4–6). The reasons why some epidemic and transmissible *P. aeruginosa* strains dominate and displace others during infection remain poorly understood, but one possibility is that intra-species competition is driven by the production of bacteriocins known as pyocins.

Pyocins are ribosomally synthesised bacteriocins which are produced to kill competitors of the same species (7). The pyocins produced by *P. aeruginosa* are classified into three major types: S, R- and F-pyocins (8). The S-pyocins are large multi-domain polypeptides which have a cognate immunity protein that binds to and inactivates the catalytic domain of the active pyocin (7). R and F-type pyocins are defective prophages ancestrally related to P2 and lambda phages respectively, which have differentiated into bacteriocins (8, 9). The R and F pyocins (collectively referred to as tailocins) possess no genetic material and therefore are nonreplicating (8, 10). Unlike the S-pyocins, the tailocins do not have a cognate immunity protein and resistance to them is mediated through incompatible lipopolysaccharide (11). Antimicrobial activity of S and R-type pyocins against planktonic cells of *P. aeruginosa* has previously been demonstrated (12–14), and pyocin S2 has been shown to be more effective against *P. aeruginosa* biofilms than aztreonam and tobramycin, both commonly prescribed antibiotics for chronic infections in CF patients (15). The killing mechanism of R-type pyocins is via a single hit membrane depolarisation, where one pyocin unit causes the death of a cell regardless of the number adsorbed to its surface (7, 8). The narrow specificity of killing has triggered interest in developing them as potent therapeutic alternatives to antibiotics (12).

Whilst some work has been performed on the role of R-pyocins in biofilm formation (16), little work has focused on whether they influence strain competition and dominance in biofilms and ultimately *in vivo*. Here we investigate whether R-pyocins are important factors in competitive interactions between *P. aeruginosa* strains taken directly from CF lungs from different patients. We show that R1 and R2 type pyocins have potent activity against established biofilm communities of sensitive *P. aeruginosa* cells. By performing pair-wise interactions between isolates, we identified two strains, A018 and A026, with reciprocated killing activity against each other. By making defined R-pyocin mutations in these strains, we determined that this activity was due to the production of different R pyocin sub-types. We found that A026 produced an R1 type pyocin which was responsible for strain displacement and domination in both planktonic cultures and multicellular biofilms. More generally, our findings demonstrate a role for R-type pyocins in shaping ecological interactions and biofilm formation in *P. aeruginosa*. We also highlight the potential of R-pyocins as antimicrobial agents against *P. aeruginosa* at a time when new antimicrobial therapies are desperately needed.

## Results

### Reciprocity of killing activity between paired *P. aeruginosa* CF isolates

We first determined whether pyocins could play a role in ecological interactions between strains isolated from CF lungs, by competing strains against each other. We isolated 24 *P. aeruginosa* CF strains from individual adult and pediatric patients, and we screened each one for pyocin killing using a spot assay to determine biological activity (17). We identified reciprocal activity between 9 strain pairs (A026/A014, A026/A018, A026/P003, A026/P013, A026/A007, A026/A024, A026/A032, A026/P010 and A014/A033), which was mediated by different subtypes of pyocin (Table S1). We found 8 pairs with one common strain partner (A026). Reciprocated killing could have been due to any of the different pyocin subtypes, therefore we investigated the class(es) of pyocins underpinning the strain antagonism that we observed.

### Reciprocal killing of paired isolates is mediated by R-pyocins

We screened for S and R-pyocin subtypes in the 24 strains using primers specific to unique regions of each pyocin gene/operon (Table 2). We used 6 primer pairs for 6 subtypes of S-pyocins (S1, S2, S3, S4, S5 and AP41) and 3 for the R-pyocins (R1, R2-group and R5). The R2 group of primers consisted of subtypes R2, R3 and R4 because they are essentially the same subtype with 98% homology in their gene sequences. The profile of the distribution pattern is shown in Table 2. We found that the distribution of S-pyocin subtypes was varied, with most strains having one or more subtype. However, 2 strains (A026 and P015), did not have any of the screened S-pyocin genes. In contrast, the presence of an R-type pyocin gene operon was uniform, with each strain having a single R-pyocin subtype. We discovered that the R1 subtype was the most represented (15 isolates), whereas the least represented was R5 (1 isolate) (Table 2). As A026 did not possess any of the tested S-pyocin genes in its genome, but was involved in reciprocal killing of other strains (Table S1), we hypothesized that the killing using the spot test assay was due to one of the tailocins (R-or F-type pyocins). We amplified an R1 pyocin sequence from A026, which was in contrast to the R2 pyocin type found in all of the 8 pair-members involved in the reciprocal interactions. This data was suggestive about the involvement of R-pyocins in reciprocal antagonism. To test this, we constructed null-R-pyocin deletion mutants (ΔR) in A026 and 4 strains (A014, A018, P003 and P013) that showed reciprocated antagonism against A026. We then repeated the spot test using the cell free extracts of the ΔR strains and showed a loss of the biological inhibition in all the 5 isolates against the protagonist strain (Fig. 1).

**Fig. 1.**
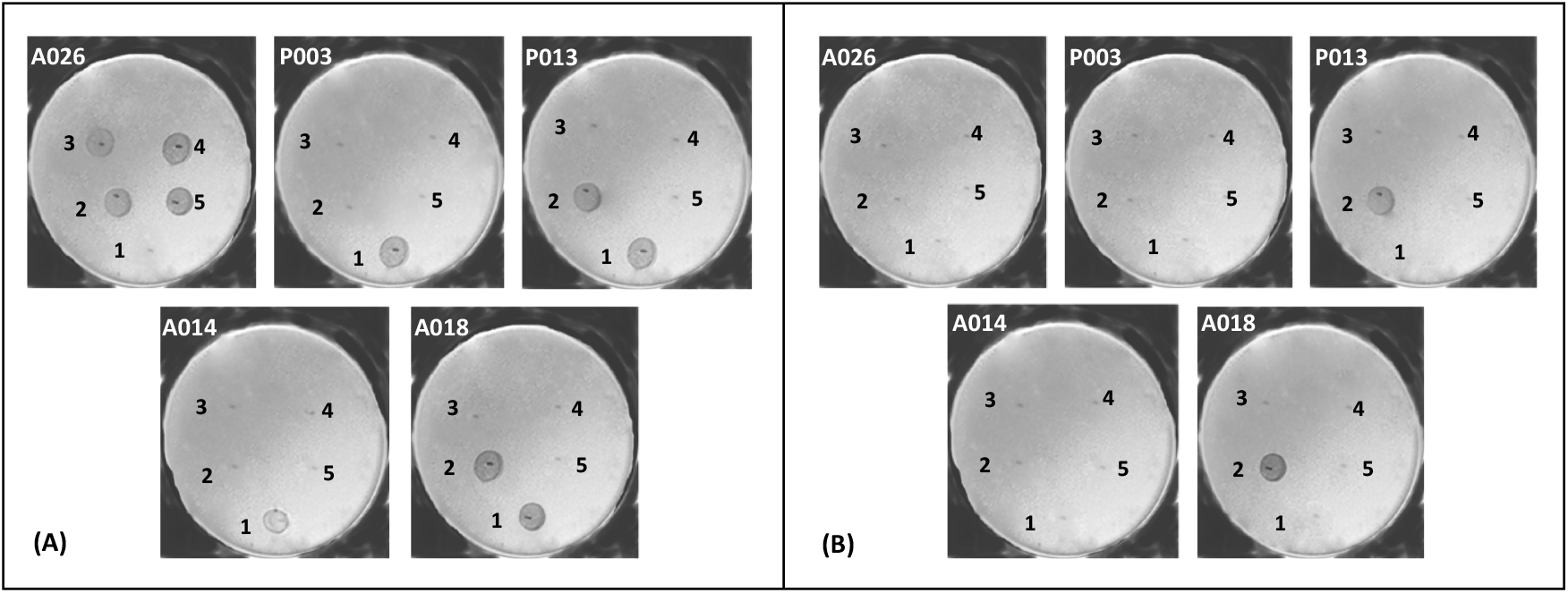
Spot test showing the biological activities of five representative clinical *P. aeruginosa* strains. Each plate is labelled with the indicator strain while numbers 1–5 represent test strains. [1= A026, 2= A014, 3= P003, 4=P013, 5= A018], In set (A), the cell-free extracts of the wild types were used for the spot assay, while set (B) used the cell-free extracts of the null-R pyocin mutant strains.

### R-pyocins mediate competition among strains existing in the same microenvironment

The role of R-pyocin in competition was previously reported in planktonic cultures of the laboratory strains PAK, PAO1 and PA14. PAK was out-competed by the other two strains, which both produce R2 pyocin, unlike PAK, which contains a mutated gene *prf6* in its R-pyocin locus, thus one competitor was weaker than the other (18). Here, we explored the involvement of R-pyocins in a new concept of head to head reciprocal killing which allows for a comparison of bi-directional pyocin activity and killing when either competing strain produces a lethal R-pyocin to the other. To complement the biological activities demonstrated in the spot assay (Fig. 1), we chose a representative competing pair A026/A018 to study in further detail. We chose A026 (R1 producer) due to its prominence as a central competitor in 8 out of the 9 pairs, and we chose A018 (R2 producer) due to the comparative demographics of the two patients from whom A018 and A026 were isolated (aged 27 and 32 respectively, with both diagnosed with CF as neonates). Both strains also had comparable growth rates which was determined by growth curves. We used transwell membrane plates to compartmentalize the two strains, while allowing cell-free media to pass through the membrane pores. We studied wild types (WT) and R-pyocin mutants (ΔR) of each strain in varying head to head combinations. Fig. 2 shows the survival rate of the WT of each strain when cultivated in adjacent wells of either the WT or ΔR derivative of the competing strain. We used sterile growth media as a control treatment. In paired interactions involving WT and ΔR strains of A026 and A018, we found that the dominant surviving strain was A026WT (Fig. 2A & B), although towards the end of the experiment, approximately 20 % of live A018WT cells persisted in the population (Fig. 2A). As R-pyocins are induced by stress, we speculate that A018 was able to persist in the population because less A018 cells reduced competition with A026, potentially leading to reduced R-pyocin production and subsequent killing by A026.

**Fig. 2.**
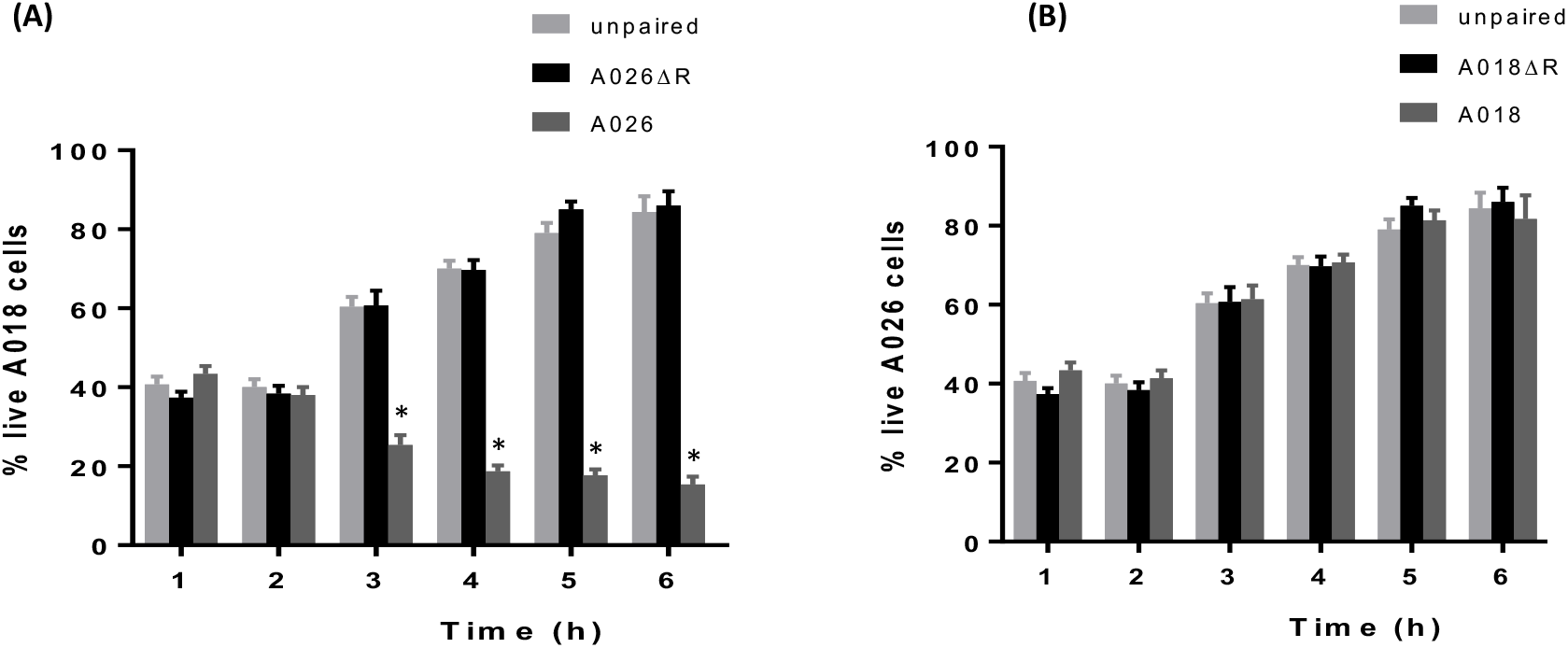
Planktonic cell competition assay in transwell plates. (A) A018 strain was grown alongside 3 different treatments (A026 wild type, A026ΔR or plain LB), and the percentage of surviving A018 cells in the population was determined over time; (B) A026 strain was grown alongside 3 different treatments (A018 wild type, A018ΔR or plain LB), and the percentage of surviving A026 cells in the population was determined over time. Hourly estimation of the percentages was achieved by staining with BacLight^®^ LIVE/DEAD stain and viewing with a confocal laser scanning microscope. Live cell counts were performed in three fields for each reading. * p value < 0.001.

### R-pyocins drive intra-species competition in biofilms

R-pyocins were shown to be important for the competition of planktonic cells in strains occupying adjacent compartmentalized environments (Fig. 2), but this does not represent the conditions found in ecosystems where strains compete side by side in biofilms for survival. In order to investigate the effect of R-pyocins on two interacting strains which co-exist in a common ecological niche, we competed strains in biofilms. We achieved this by mixing A018 and A026 differentially labelled with GFP or mCherry in microfluidic channels of a BioFlux® device and allowed them to form biofilms over time. In total we used six combinations of labelled strains (Fig. 3). When A026WT labelled with either GFP (A026_GFP_) or mCherry (A026_m-Cherry_) was incubated with A018WT labelled with the contrasting fluorophore, we found that A026 always outcompeted A018 (Fig. 3). However, when A018 WT was mixed with A026ΔR, A018 outcompeted A026ΔR (Fig. 3). When R mutants of both strains (A018ΔR and A026ΔR) were competed against each other, both strains were able to co-exist within biofilms, but they were clearly spatially separated, enabling each to form individual pockets within biofilms which were strain specific (Figs. 4A & B). Since inactivating R-pyocins in strains with different R-pyocin types abolished competition, we explored whether 2 strains with matching R-pyocin types could co-exist. We mixed 2 strains, A018 and P013, which were tolerant of each other in the spot test assay (Table S1) and possessed the same R and S-type pyocins (Table 2), to study the patterns of biofilm formation. We found that both strains could co-exist and produced biofilms with a similar architecture to the R-mutant biofilms of A018 and A026 (Fig. 4C).

**Fig. 3.**
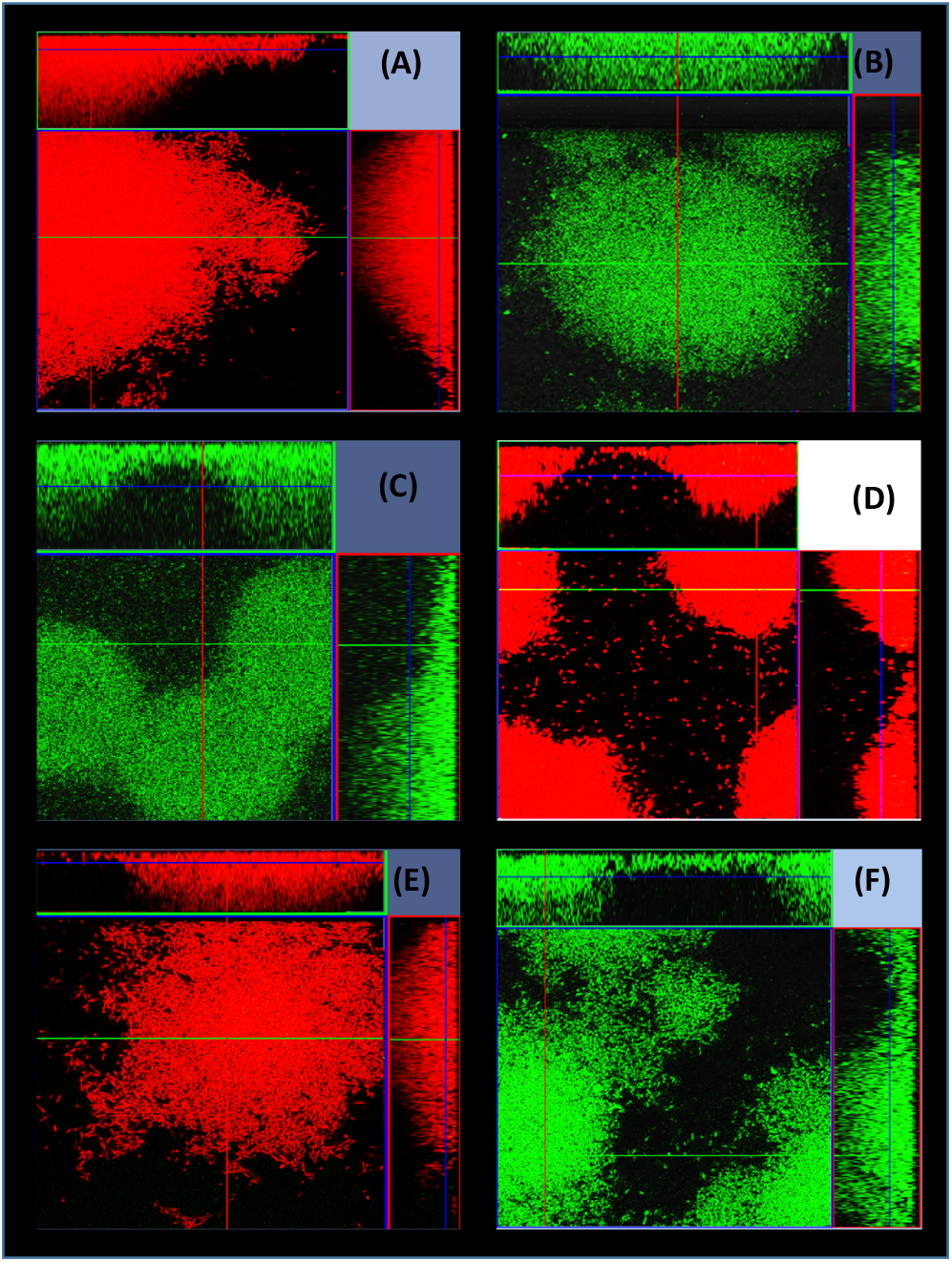
Strain competition in biofilms. 15 h biofilms developed in a microfluidic BioFlux® device as a result of competition between A026 and A018. A026 dominated in the biofilm competition of A018 vs A026 and A018ΔR vs A026 regardless of the fluorophore (mCherry or GFP) used to tag either strain – Figures A-D. However, A018 dominated when in competition with the null-R-mutant of A026 (A026ΔR) – Figures E-F. Key: (A) =A026mC vs A018GFP, (B) = A026GFP vs A018mC, (C) = A026GFP vs A018ΔR-mC, (D) = A026mC vs A018ΔR-GFP, (E) = A026ΔR-GFP vs A018mC, (F) = A026ΔR-mC vs A018GFP

**Fig. 4.**
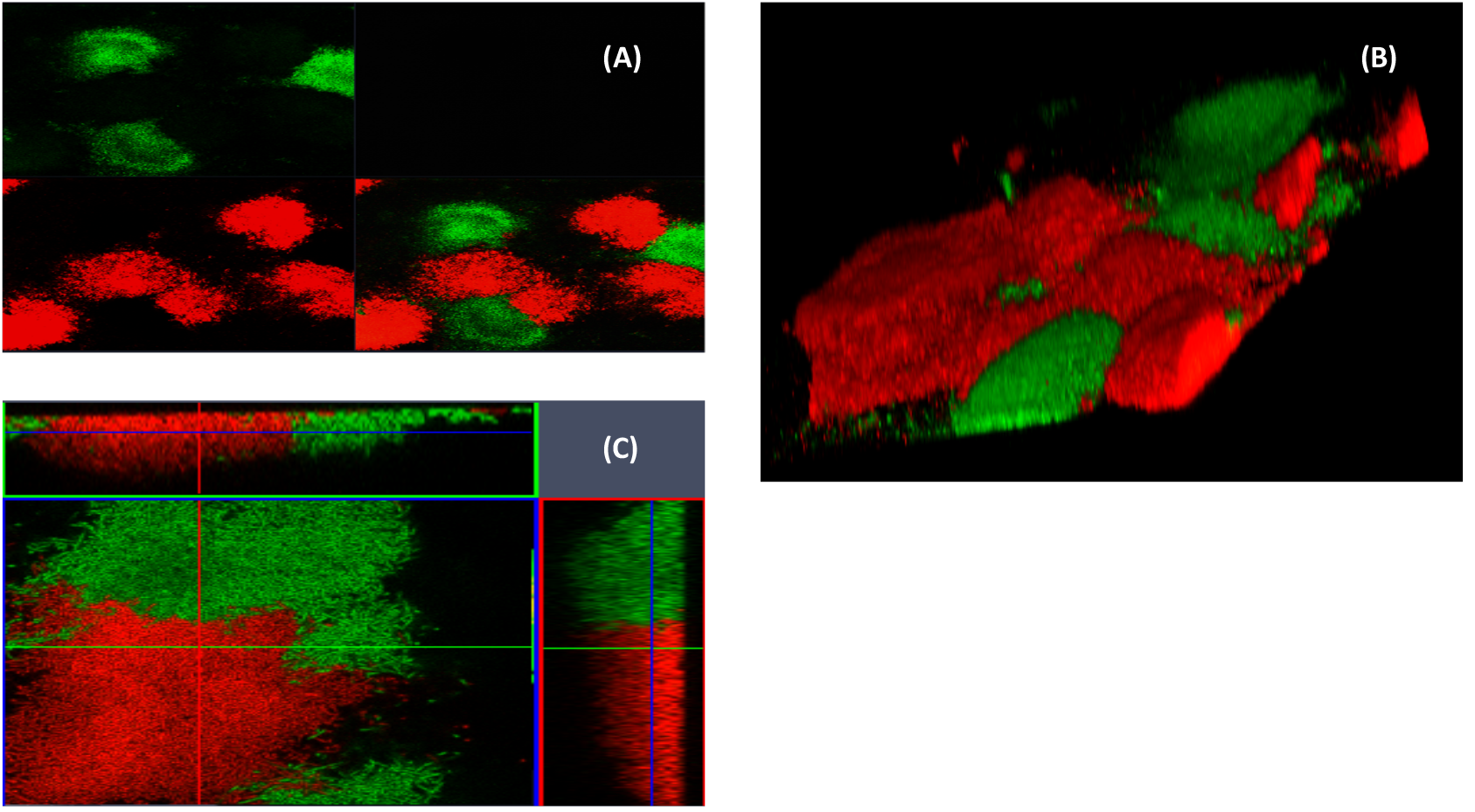
15 h biofilms showing the distribution of two strains. A and B show biofilms of mixed cultures of A018ΔR-GFP and A026ΔR-mCherry. (A) is a split image of the fluorescence green and red channels at 10x magnification showing a wider distribution of the strains as they form micro-niches whilst (B) shows a closer view of the same biofilms at 63x magnification. (C) shows a biofilm of wild types A018GFP and P013mCherry which both have the same subtypes of S- and R-pyocins.

### R-pyocins demonstrate anti-biofilm properties in established biofilms

The loss of biological activity in the R-pyocin mutant spot test suggested the involvement of this class of pyocin in biofilm killing activity. We induced R-pyocin production in either A018 or A026 to purify high pyocin yields and tested these against biofilms. We subjected biofilms of A018 or A026 grown on polypropylene beads, to a single treatment of R-pyocins from the competitor strain for one hour and we counted viable cells before and after treatment over time. We found that R-pyocins of each strain caused a significant reduction of viable cell counts of the competing strain (Fig. 5A). However, extracts from the R-pyocin mutants of either strain did not produce any significant differences in the viable cell count of the competitor when compared to the untreated population (Fig. 5B). We also used a BioFlux system to achieve a longer exposure time to R-pyocins and a dynamic flow. We found that mature biofilms treated with R-pyocins, resulted in live cell populations diminishing significantly over time, with a corresponding increase in the number of dead cells. There was a progressive loss in the number of viable cells in biofilms and full thickness biomass eradication was achieved within 4 h of starting R pyocin treatment using purified R-pyocins of wild type strains (Fig. 6A shows antibiofilm activity of A026 R-pyocin against A018 biofilm). In contrast, the extract purified from the R-pyocin mutants, showed no adverse effect on the biofilm growth and maturation of the competing strain (Fig. 6B shows A026ΔR extract against A018 biofilm). The results were similar when A026 biofilms were treated with the R-pyocins/ extracts from A018 and A018ΔR respectively.

**Fig. 5.**
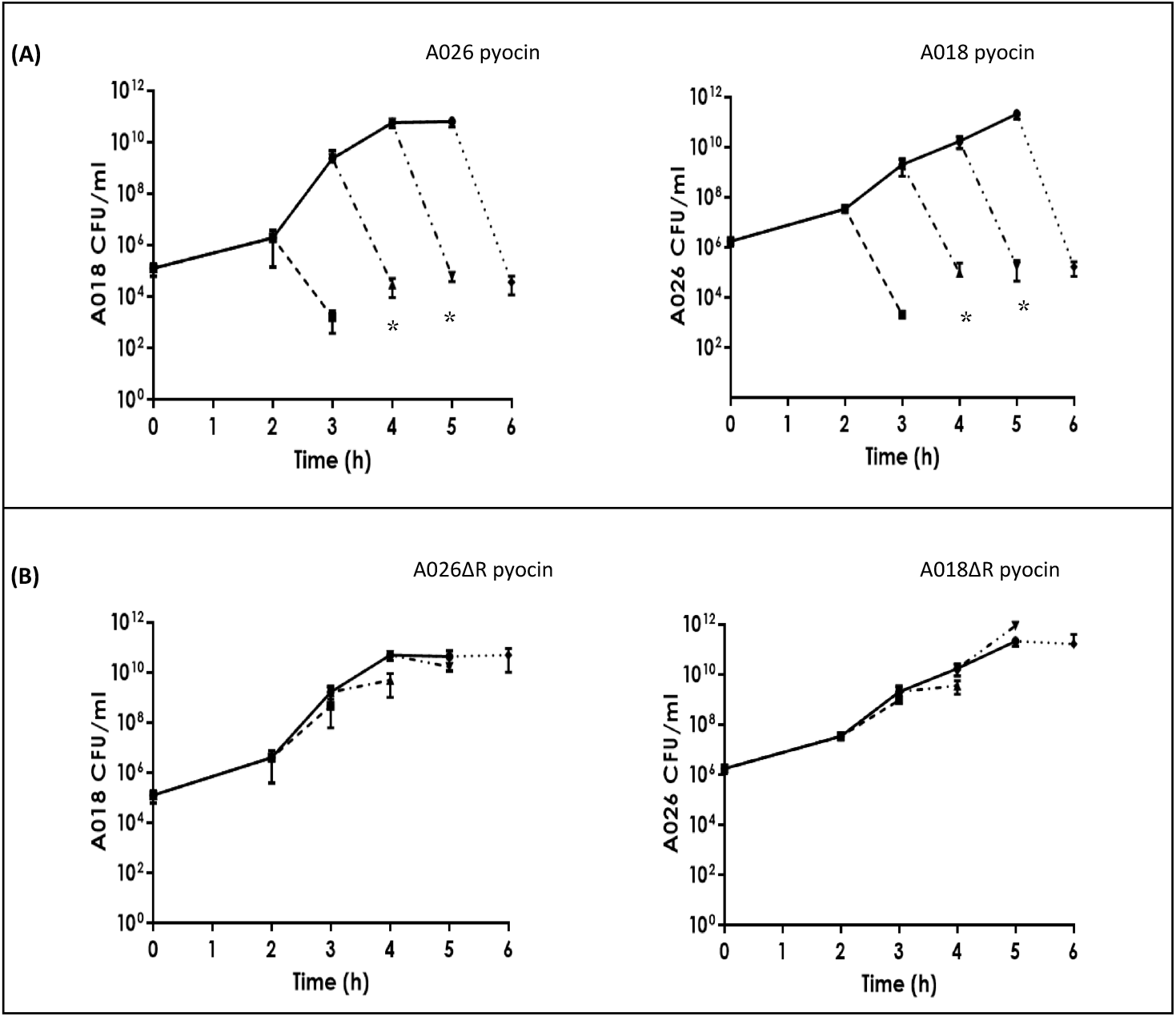
Single static treatment of biofilms of A018 or A026 grown on polystyrene beads using R-pyocins of a competitor (A026 or A018 respectively). Hour 0 readings are the CFU/ml values of the 24 h biofilm grown on the beads. In one set of beads, growth was allowed to proceed unhindered (unbroken lines) while in the other set, three beads were harvested every hour, their biofilms were treated for one hour using purified R-pyocins and the CFU count was recorded after treatment (broken lines). Beads in (A) were treated with R-pyocins from wild type of competitor while the ones in (B) were treated with cell-free extracts from R-pyocin mutants (* p < 0.05).

**Fig. 6.**
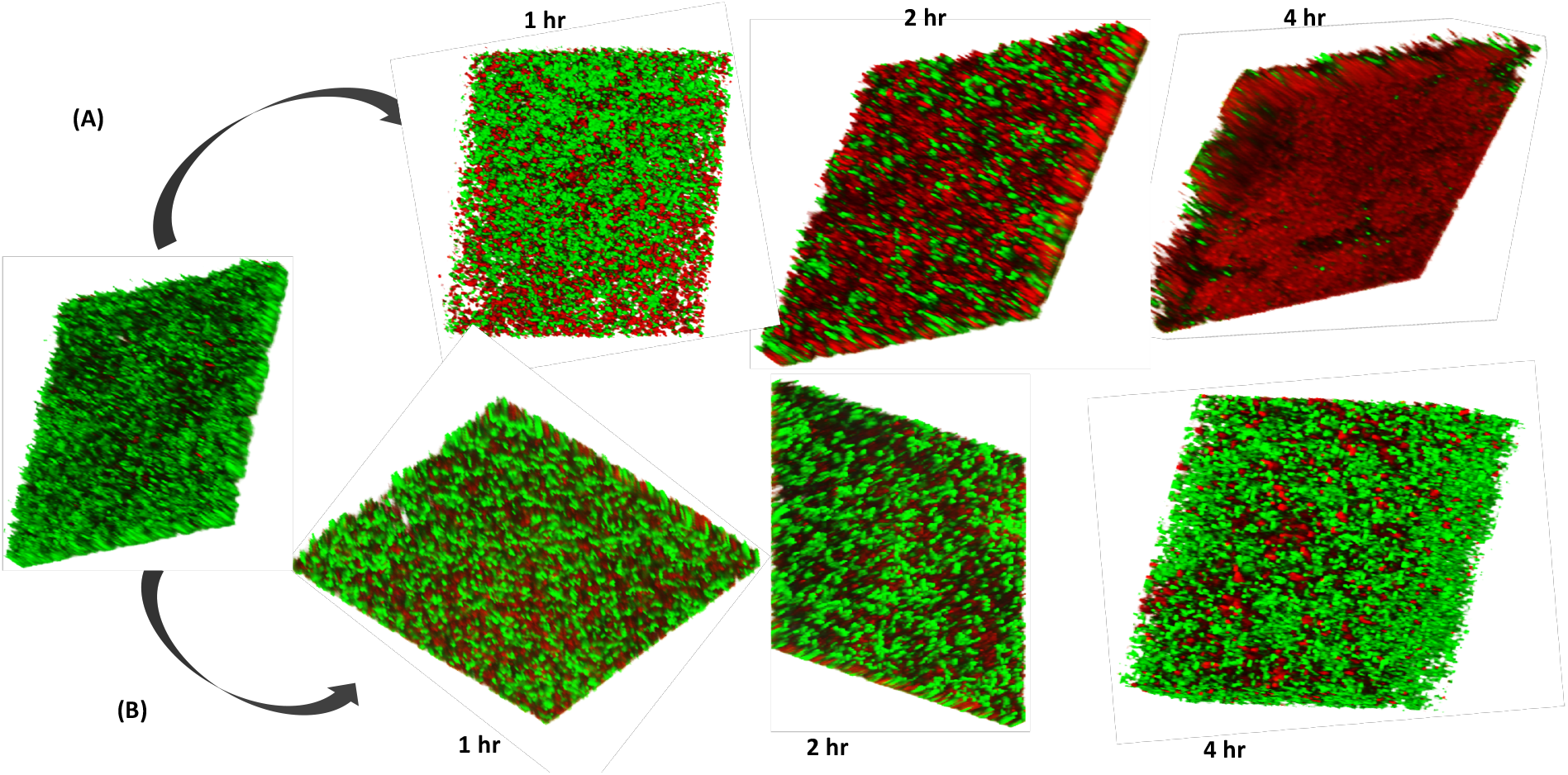
Anti-biofilm efficacy of R-pyocins. A 15 h biofilm of A018 was treated with R-pyocins extracted from A026 (A) and A026ΔR (B). A significant portion of the biomass was killed after 2 h and full-depth lethal effects on the biomass was achieved after 4 h. This effect was absent in the control experiment which utilised the R-pyocins of A026ΔR.

## Discussion

A key clinical problem and research priority in CF is how to treat and eradicate *P. aeruginosa* from the CF lung, and how to prevent pulmonary exacerbations (3, 19). This has proven incredibly challenging and is likely due to the overwhelming phenotypic and genomic diversity that we now know evolves within *in vivo P. aeruginosa* populations (4–6). Some epidemic strains, such as the Liverpool Epidemic (LES) or Denmark epidemic (DK2) strains, have become extremely successful, infecting many different patients across multiple geographic locations. How these transmissible and persistent *P. aeruginosa* lineages displace and prevent the colonization of other strains remains poorly understood, but elimination of competing strains is one potential survival strategy. Competing against genetically similar strains is an important means of self-preservation, because available nutrients and space in an ecological niche are more likely to be competed for between highly-related strains.

Certain factors have previously been suggested to influence strain succession and persistence. Production of a compound from environmental pseudomonads possessing a homologue of the thioquinolobactin gene cluster was attributed to strong antagonistic effects on clinical isolates of *P. aeruginosa* (20); while the role of temperate phages in improving competitive fitness in CF lungs was recently reported (21). In addition, over 90% of known strains of *P. aeruginosa* are able to produce pyocins (7, 8). These bacteriocins often possess a narrow spectrum of antimicrobial activity against closely related strains and so are likely to be involved in interstrain competition. Recently it has been shown that strains of *P. aeruginosa* isolated from CF lungs are particularly susceptible to R-pyocins, mainly due to their LPS architecture with non-typeable or non-accessible O-serotypes (22).

In this study, we provide support that R-type pyocins help shape ecological interactions between CF isolates of *P. aeruginosa* in biofilms, and that R-pyocins have potential as antimicrobial agents. Specifically, we found that (i) individual *P. aeruginosa* strains taken from different CF patients produced only one R-pyocin type, whereas the number of S-pyocin types produced by any one strain varied; (ii) R-pyocins were crucial for strain dominance during competition assays in planktonic cultures and within biofilms, and where variations in both R and S subtypes occurred in the competing strains; (iii) purified R-pyocins from different strains demonstrated significant antimicrobial activity against established biofilms.

We began our study by isolating 24 *P. aeruginosa* strains from 24 different CF patients, and we identified several isolates with reciprocated antagonism in pairwise interactions. Deletion mutagenesis of essential regions of R pyocin genes of each strain, revealed that the antagonistic activity was due to the production of R-pyocins. While the number of S-subtype pyocins differed per strain, ranging from zero to all six tested S type pyocins (Table 2), each strain had only one R-subtype. With little genetic redundancy in the bacterial genome, the incorporation of an extra S pyocin gene will be more amenable to genetic restructuring than the assembly of multiple R-type operons. Indeed, many *P. aeruginosa* strains possess extra ‘orphan’ immunity genes to the S-type pyocins presumably creating increased pyocin resistance with little extra costs (23). The diversity of S-type pyocins displayed by some of the clinical isolates did not proportionally translate to better survival in the face of competition. Contrariwise, the strain A026, which demonstrated the highest competitive ability, did not have any of the six screened S-pyocin types (Table 2).

The abolishment of killing activity in some of the clinical isolates was only achieved in R-pyocin mutants, even if a strain had an excess of S-pyocin genes, highlighting a role of R-pyocins in strain survival (Fig. 1). However, it is important to note that killing patterns between isolates using the spot assay (Fig. 1, Table 2 and Table S1), did not always show a clear and obvious role for R-pyocin killing. For example, A031 (R1 producer) was highly sensitive to pyocin extracts from other strains but also killed other R1 producers (Table S1). This suggests that A031 kills in an R-pyocin independent manner, and indeed, F-pyocins and/or phage are also likely to be important in *P. aeruginosa* strain competition. Strains can also become sensitive to their own R-pyocins through altered LPS (23), and this further complicates the picture. More work is required to unravel all the factors which drive interactions between *P. aeruginosa* strains.

R-pyocins have previously been shown to confer a growth advantage to producing strains *in vitro* during competition of planktonic cells (11, 18, 24), and promote antimicrobial activity against *P. aeruginosa in vivo* using murine models (12). However, any proposed role in strain competition *in vivo*, or the applicability of R-pyocins as potential antimicrobials, hinges on their ability to combat biofilms, the preferred mode of *P. aeruginosa* growth in chronic infections such as in the CF lung and chronic wounds. We demonstrated a direct association of R-pyocins with anti-biofilm efficacy. Although a previous study showed that sub-inhibitory concentrations of cell-free culture supernatants produced an enhancement of biofilm formation of susceptible strains (16), we found that exposing two competing strains to each other ensured R-pyocin-dependent killing in biofilms over time.

Our study also highlights the potential of using R-pyocins as an anti-biofilm strategy, and that R-pyocins can effectively eradicate biofilms composed of a susceptible strain (Fig. 6). Exposure of live cells, planktonic or biofilm, to extracted R-pyocins (Figs. 1 and 6 respectively), results in killing in the absence of a live competing strain (Fig. 2A). Therefore, R-pyocins could be a useful alternative for antibiotics or phage therapy for infections involving biofilms, or for biofilms growing on medical devices such as catheters. In contrast to phages, the absence of a capsid containing genetic material in the structure of the R-pyocins nullifies their ability for self-replication and they also have highly specific killing action (8).

In summary, R-pyocins are a class of tailocins which are likely to be important in shaping *P. aeruginosa* populations during chronic biofilm-driven infections such as those found in the CF lung and in chronic wounds. Mutation of the R-pyocin locus was a major factor that eliminated competition between strains in our study, and co-existence of strains was achieved when biofilms of R-pyocin mutants of competing strains were cultivated together. They therefore could have particular relevance in niches where competition is central to survival. More work is required in the future as to the mechanisms that regulate R-pyocin production and their role in shaping *P. aeruginosa* ecology and their potential role as therapeutic agents.

## Materials and Methods

### Bacterial strains, plasmids, media and culture conditions

We used 24 *P. aeruginosa* isolates taken from CF sputum samples (14 adult and 10 pediatric patients), and *Escherichia coli* (S17 −1 (25) and DH5α (26)) strains in this study (Table 1). We routinely cultured our strains in Lysogeny broth (LB) agar at 37 °C and cultivated the biofilms of our *P. aeruginosa* clinical strains in minimal media M9 broth supplemented with 0.2% citrate. (M9 broth is made up of basal salt (68 g/L Na_2_HPO_4_, 30 g/L KH_2_PO_4_ and 5 g/L NaCl), 10^−2^ M NH_4_Cl, 10^−4^ M CaCl_2_, 10^−3^ M MgSO_4_.7H_2_O). Using tetracycline (Tc) 25 μg/ml, we maintained and/or selected our plasmids in *E. coli* S17-1 while our *P. aeruginosa* trans-conjugants were selected on Tc 150 μg/ml + Nalidixic acid 15 μg/ml plates.

**Table 1.**
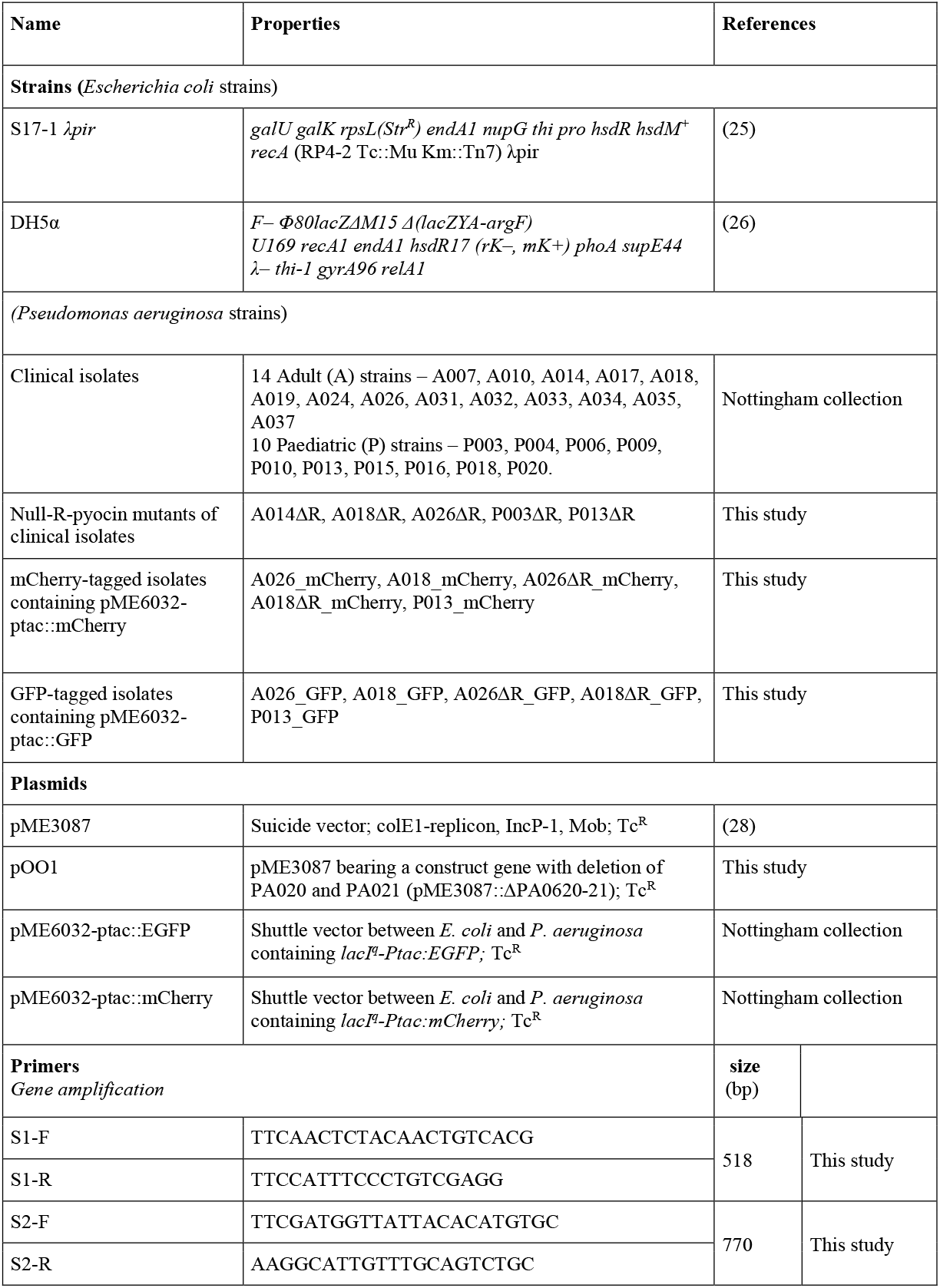
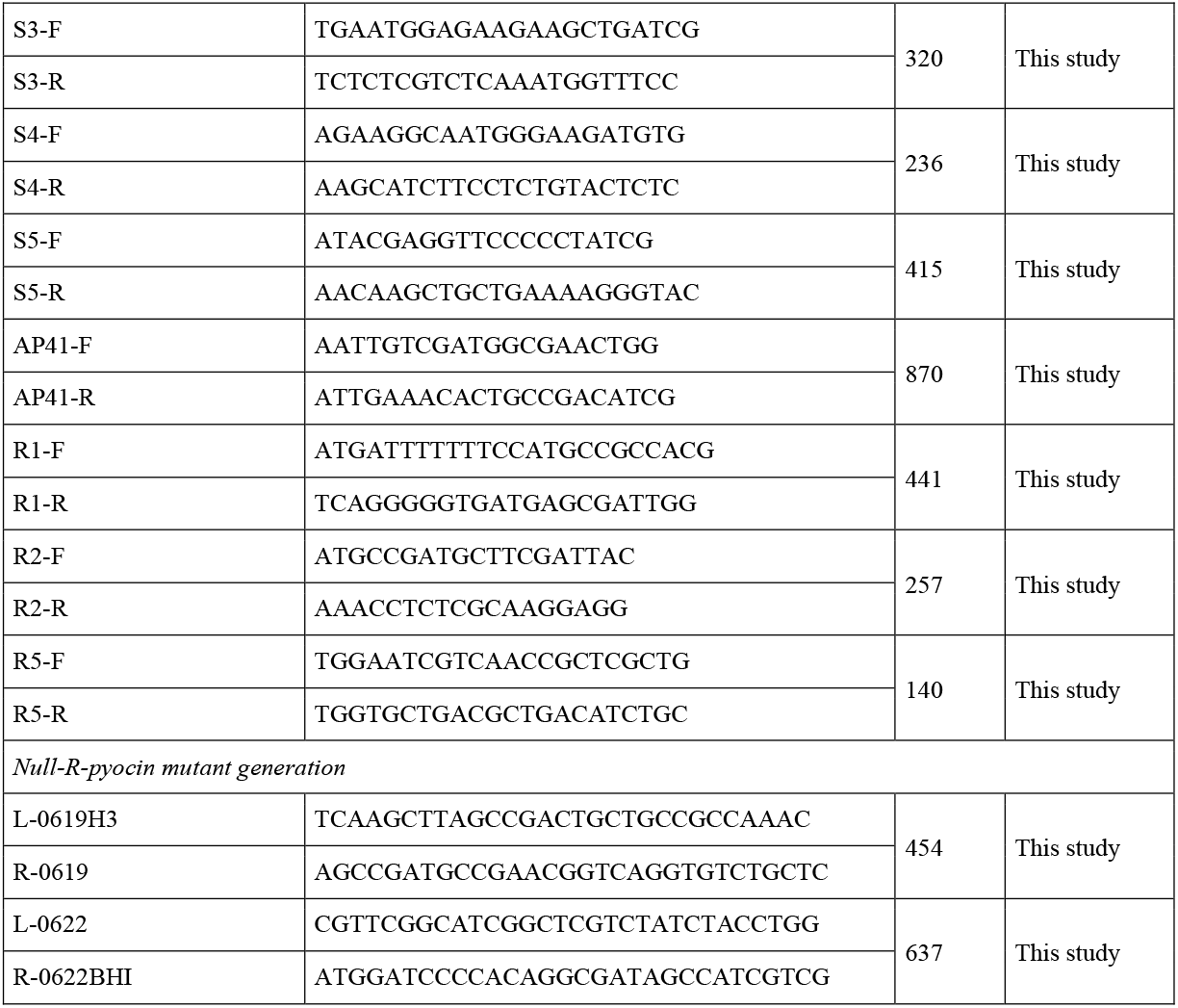
Strains (wild type and mutant derivatives), plasmids and primers used in this study.

**Table 2.**
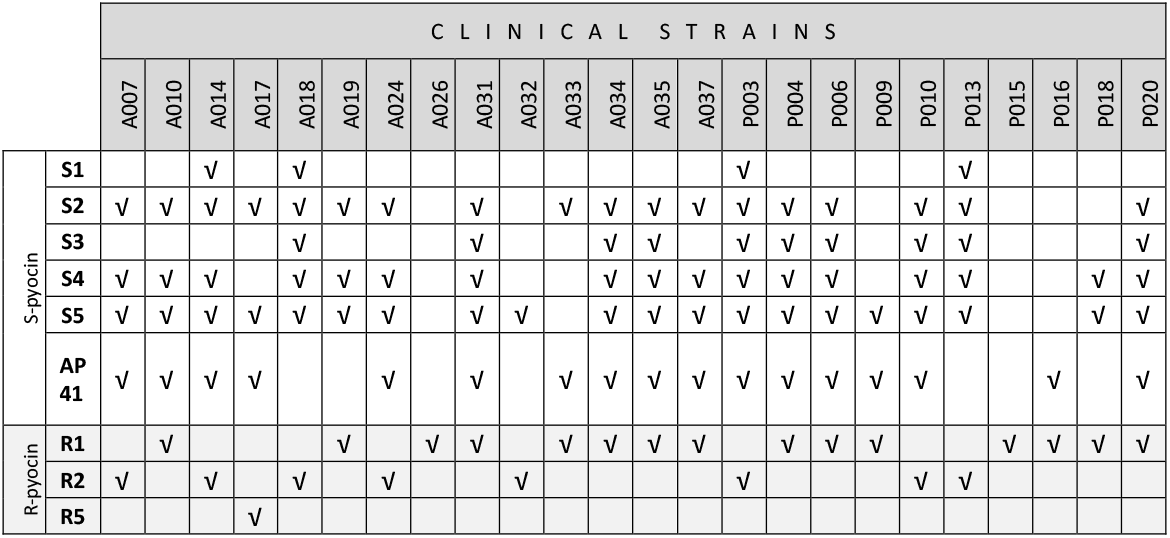
The distribution of the six S-pyocin subtypes and three R-pyocin subtypes in the *P. aeruginosa* clinical isolates used in this study.

### Polymerase Chain Reaction (PCR) conditions

All primers used in this study were designed by using the *P. aeruginosa* genome database as reference (http://www.pseudomonas.com) (27) and are listed in Table 1. Our PCR volume of 50 μl comprised 2.5 μl each of the primers (10 mM) and the template, 10 μl each of buffer and GC enhancer, 1 μl of dNTPs (10 mM), 0.5 μl of Q5 high fidelity polymerase and 21 μl of de-ionised water. The PCR conditions we used were 98 °C for 30 s, 30 cycles of 98 °C for 10 s, 58 °C for 30 s, 72 °C for 30 s/kilobase followed by a final extension at 72 °C for 2 min

### Spot assay for pyocin activity

Using 2 μl of the overnight cultures of our *P. aeruginosa* isolates (adjusted to OD_600_ = 0.5 ± 0.02), we inoculated 5 ml of cooled soft top (0.4%) agar. We poured this mixture onto LB agar plates to produce an overlay of the indicator strain. Next, we vortexed equal volumes of the overnight broths of the test strains and chloroform. We centrifuged the vortexed mixture to obtain cell free extracts from the supernatant. We spotted 7 μl drops from the supernatant of individual test strains, on the agar overlay indicator strains, allowing the spots to dry and incubating at 37 °C overnight. Clear zones of growth inhibition indicate pyocin-dependent lysis of the susceptible strains.

### R-pyocin mutant generation

We generated R-pyocin mutants by deleting the genes coding for the R-pyocin tail fibre and its chaperone assembly gene (PA0620 and PA0621 in PAO1 reference strain). We achieved this using a double homologous recombination method with the help of our deletion gene construct. Our 1076 bp gene construct was generated from a 2-stage PCR. In the first stage PCR, we amplified the upstream and downstream genes PA0619 and PA0622 (flanking our genes of interest PA0620 and PA0621) separately using primer pairs L0619H3/R0619 and L0622/R0622BHI. This first stage generated 454 bp and 637 bp amplicons respectively. In the second stage, we used the two amplicons from the first stage as templates, bringing the engineered 15bp complementary ends together by splicing of overlapping extensions using the end-primers L0619H3 and R0622BHI. Next, we cloned the gene construct into *HindIII/BamHI* digested pME3087 (28) to produce plasmid pOO1 which was used to transform *E. coli* S17-1 strains. We then integrated pOO1 into *P. aeruginosa* chromosomes by conjugating *P. aeruginosa* strains with transformed S17-1 cells. We selected our *P. aeruginosa* trans-conjugants using tetracycline 150 μg/ml + Nalidixic acid 15 μg/ml agar plates. We picked individual colonies from our selective plates and cultured them in overnight broths without antibiotics in order to enrich for tetracycline sensitive cells. We then made fresh cultures (1:100 dilution) from the overnight broth and incubated till OD_600_ = 0.1 ± 0.05. We added tetracycline and carbenicillin in turn at 1 h intervals to achieve final concentrations of 20 μg/ml and 2000 μg/ml respectively. Then, we washed our cells using sterile LB broth and repeated the enrichment process (in plain and antibiotic broths). After washing the cells, we plated out their dilutions (10^−5^ to 10^−8^) on LB agar to harvest individual colonies. We performed a replica plating of individual colonies using LB and Tc 25 μg/ml agar plates. This step selected for unmarked mutants with double cross overs and being sensitive to tetracycline, we picked them from the LB plates. We further confirmed the deletion mutation by PCR using primer pair L0619H3 and R0622BHI, thus recovering our 1076bp deletion gene construct. We tested for the successful attenuation of pyocin activity by performing spot assays using cell-free extracts from these null-R-pyocin mutants and comparing the results with cell-free extracts from wild-type bacteria.

### Induction and purification of R-pyocins from *P. aeruginosa*

This is a modified method adopted from (29). Briefly, starting with 1:100 starter culture from an overnight *P. aeruginosa* broth, we achieved log phase (OD_600_ ~0.25) in 2 L of G-medium (20 g/L sodium glutamate, 5 g/L glucose, 2.23 g/L Na_2_HPO_4_, 500 mg/L yeast extract, 250 mg/L KH_2_PO_4_ and 100 mg/L MgSO_4_.7H_2_O). We then added mitomycin C to a final concentration of 3 μg/ml. After incubating the culture for a further 3 h; we removed the cell debris by centrifuging at 17,000 x *g* for 1 h at 4°C. To the supernatant, we added 4 M (NH_4_)_2_SO_4_ titrating this at a rate of 1 ml/min with continuous stirring at 4 °C. After an overnight storage at 4 °C, we centrifuged the suspension at 22,000 x *g* for 1 h to harvest the pellets containing the R-pyocins. We resuspended the pellets in TN50 buffer (50 mM NaCl and 10 mM Tris HCl adjusted to pH 7.5) and further performed ultracentrifugation at 65,000 x *g* and 4 °C for 1 h to further concentrate the R-pyocins. We resuspended and stored our R-pyocin rich pellets in 2 ml TN50 buffer at 4°C.

### Transwell membrane plate method for planktonic cell activity

We used a standard 6-well plate with a transwell polyester membrane (Corning®) to study cell-free interactions between *P. aeruginosa* isolates placed on either side of the membrane. The transwell units are made up of coupled multi-well plates. Each couple has an outer well set up in a plate of six wells and individual insert wells. The base of the insert has a permeable support through which cell-free exchange of culture media take place. The membrane pores are 0.4μm in diameter through which soluble and particulate pyocins (R-pyocin dimension 120nm by 5nm, (30)) are freely exchanged. Cultures of the two strains are grown separately, adjusted to comparable optical densities and placed in separate wells of each transwell couple. We started the initial cultures of either isolate at an OD_600_ value 0.05 ± 0.01 with each pair set up in triplicates using 2.5 ml volume in each well. We incubated the culture plates with gentle shaking (60 rpm) at 37 °C for 2 h. Thereafter, we took 100 μl samples from either well and washed twice with phosphate buffered saline (PBS). We then stained the cells using Baclight® LIVE/DEAD stain by incubating them in the stain for 30 min at 37°C. We used this staining method for each cell population on either side of the transwell membrane and viewed them under the confocal microscope. We carried out two sets of confocal studies. In the first set, we viewed the cells in their native state with minimal disturbance by skipping the PBS washing step. The aim of the second set was cell counting, so we washed and diluted out the cultured cell populations. We counted three fields for each cell culture at 60 x magnification. We included two controls, these were null-R pyocin mutants of either strain paired up with wild type of the competitor and unpaired i.e. free growing wild type of either strain with plain LB medium in the adjacent well. Using the live and the total cell counts, we calculated the percentage live cells in different fields over time.

### Biofilm development on 3D pearl beads and pyocin treatment

We used minimal media M9 broth for our bead biofilm cultivation. These beads, in hollow cylinder shapes, were made of polystyrene and had an approximate total surface area of 2.6 cm^2^. Starting with 1:100 dilution of culture, we suspended these sterile beads in 100 ml of fresh M9 culture in a 500-ml flask. The incubation condition we employed were 37 °C, shaking at 80 rpm for 24 h. After 24 h, we harvested the beads and re-suspended them in fresh broth that had been grown to log phase. At this stage, we started the timing of the experiment, taking the start of the new cultures as 0 h. At 2 h, we commenced hourly harvesting of six beads from the lot. The harvested beads were gently rinsed twice in PBS to rid them of planktonic cells. We divided these six beads into two sets of three. We used the first set for CFU cell counting by vortexing them in 1 ml PBS and plating them in dilutions. We used the second set to study R-pyocin treatment by suspending the beads in purified R-pyocins for 1 h and thereafter performing the cell count. Our controls included a third set of beads that were treated with cell-free extracts from null-R mutant derivatives of each isolate.

### Growing mono and mixed culture biofilms in a microfluidic system

We employed the microfluidic system of BioFlux 200 (Fluxion Biosciences Inc., CA, USA) to cultivate our biofilms. The system comprised twin wells (inflow and outflow) connected by microchannels for cultivating biofilms. As an initial step, we primed the channels with 100 μl of LB medium from the inflow well with a share pressure setting of 2 dyne/cm^2^ for 3 min. Thereafter, we seeded the microchannels from the outflow well using fresh cultures at mid-log growth phase (approx. OD_600_ = 0.05) by applying a back pressure of 0.5 dyne/cm^2^ for 2 s. We then incubated the set-up on the heating plate at 37 °C for 30 min without flow to enhance adhesion of the cells to the channels. We coupled the microfluidic system for overnight incubation (approx. 15 h) with pressure and temperature settings of 0.25 dyne/cm^2^ and 37 °C respectively. The strains that we used to seed the microchannels and cultivate our biofilms were previously tagged with either green fluorescence protein (GFP) or mCherry. Our biofilms were cultivated as either mono-or mixed cultures. Our mono-cultures consisted of A026_GFP_, A026_m-Cherry_, A018_GFP_, A018_mCherry_, A026ΔR_GFP_, A026ΔR_m-Cherry_, A018ΔR_GFP_, A018ΔR_m-Cherry_, P013_GFP_, P013_m-Cherry_, P013ΔR_GFP_ and P013ΔR_m-Cherry_; while the mixed cultures consisted of differentially labelled pairs i.e. A026_m-Cherry_/A018_GFP_, A026_GFP_/A018_m-Cherry_, A026_GFP_/A018ΔR_m-Cherry_, A026ΔR_m-Cherry_/A018ΔR_GFP_ and A018_m-Cherry_/P013_GFP_ among others. After 15 h, we viewed our biofilms using a confocal laser scanning microscope (Zeiss® LSM 700).

### R-pyocin treatment of biofilms in a microfluidic system

We studied a dynamic flow of pyocin treatment on biofilms cultured in the BioFlux 200 microfluidic system. To achieve this, we cultivated 15 h biofilms, staining the cells as they multiplied, by adding the Baclight LIVE/DEAD stain to the 10% LB broth used to cultivate the biofilm. We then treated the 15 h mature biofilms with purified R-pyocins by changing the medium to the TN50 buffered R-pyocin suspension mixed with the LIVE/DEAD stain. We maintained the same treatment conditions (temperature and pressure) for a further 5 h. We included two controls in the study, these were biofilms treated with null-R mutant extracts and untreated biofilms.

## ACKNOWLEDGEMENTS

We thank the University of Nottingham for the award of a Vice-Chancellor’s Ph.D scholarship to OO.

---

**Table S1.**
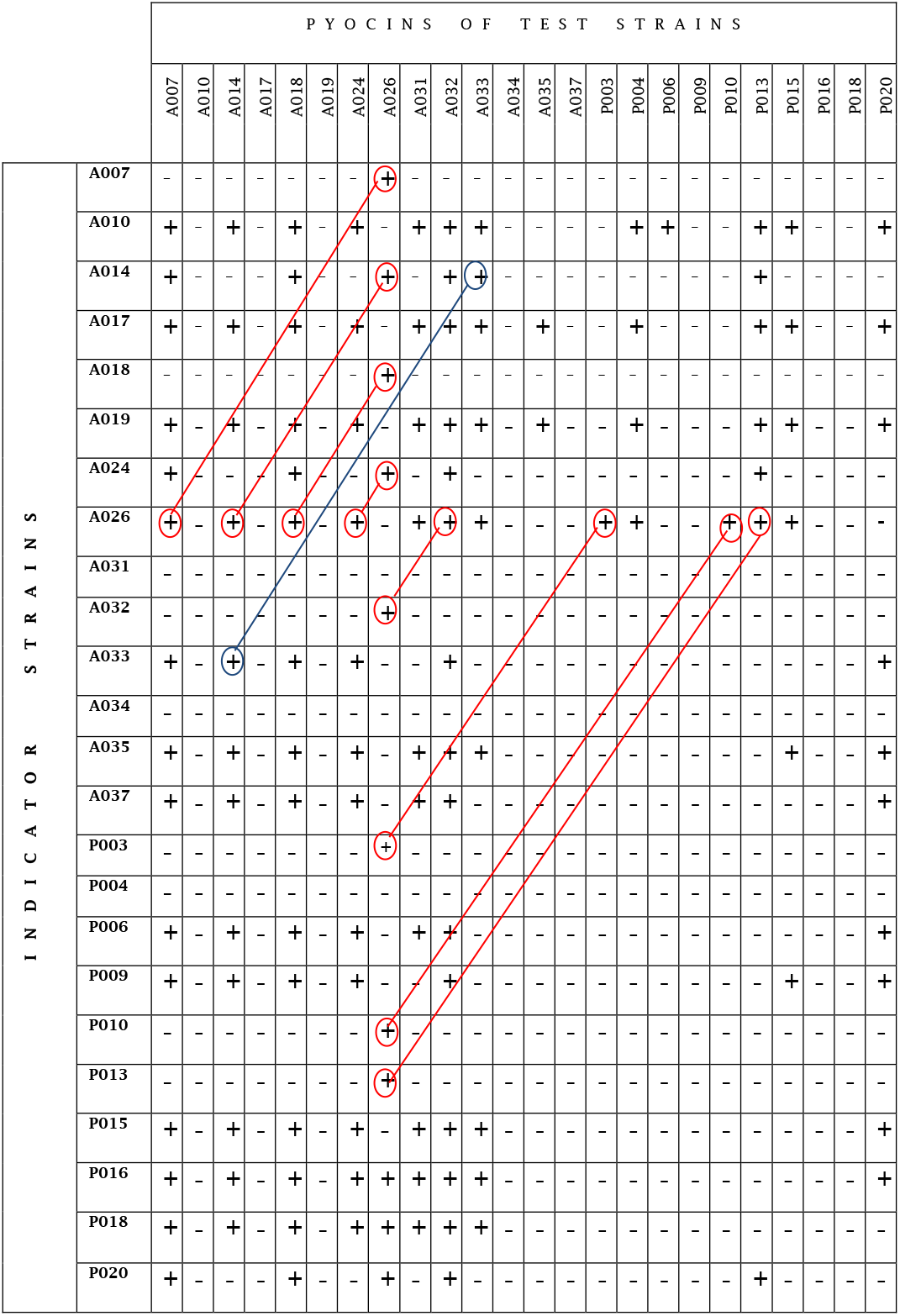
Interactions of 24 clinical *P. aeruginosa* strains using a spot test assay of biological activity. Vertical columns show the pyocin activities of each test strain while the rows show each isolate as an indicator strain. Pairwise strain antagonism are represented as dumb bells. Red dumb bells for pairwise activities having A026 as a competing member while the blue dumb bell represents reciprocal killing between A033 and A014 (+ = lethal activity shown by zone clearance, − = no activity).

Author contributions
O.O., C.N.P. and S.P.D. conceived the study, analyzed data and wrote the paper; O.O. performed the experimental work.

